# Exploratory methods for high-performance EEG speech decoding

**DOI:** 10.1101/2021.11.16.468876

**Authors:** Lindy Comstock, Claudia Lainscsek, Vinícius R. Carvalho, Eduardo M. A. M. Mendes, Aria Fallah, Terrence J. Sejnowski

## Abstract

State-of-the-art technologies in neural speech decoding utilize data collected from microwires or microarrays implanted directly into the cerebral cortex. Yet as a tool accessible only to individuals with implanted electrodes, speech decoding from devices of this nature is severely limited in its implementation, and cannot be considered a viable solution for widespread application. Speech decoding from non-invasive EEG signals can achieve relatively high accuracy (70-80%), but only from very small classification tasks, with more complex tasks typically yielding a limited (20-50%) classification accuracy. We propose a novel combination of technologies in which transcranial magnetic stimulation (TMS) is first applied to augment the neural signals of interest, producing a greater signal-to-noise ratio in the EEG data. Next, delay differential analysis (DDA) – a cutting-edge computational method based on nonlinear dynamics – is implemented to capture the widest range of information available in the neural signal, by incorporating both linear and nonlinear dynamics.

## I. INTRODUCTION

In recent years, the field of speech decoding has made significant advances that place the goal of real-time translation from neural signals within sight [28, 44]. At the same time, current technologies still suffer from major limitations that require innovative new solutions to overcome. Brain-computer interface (BCI) devices that utilize neural signals from invasive implants remain inaccessible for the majority of their target user population [40], whereas BCI devices that decode from electroencephalography (EEG) signals remain insufficiently fast with low accuracy and a limited range of classification abilities [32, 35, 36]. Ultimately, the most realistic chance for wide-spread application of speech decoding technology will require a non-invasive method that can achieve a similar classification accuracy accomplished by invasive methods. This pilot study aims to increase the accuracy and generalizability of speech decoding from non-invasive EEG neural signals by means of a novel combination of technologies: the application of transcranial magnetic stimulation (TMS) to augment the neural signal of interest, resulting in greater signal-to-noise ratio in the EEG data, and delay differential analysis (DDA), a cutting-edge computational method that incorporates both linear and nonlinear dynamics to capture the widest range of information available in the neural signal.

Currently, the best metrics for speech decoding from EEG signals are obtained by means of a convolutional neural network (CNN) [36]. However, these results only achieve high performance (78-89%) for simple binary classification tasks which register the presence or absence of certain categories of articulatory movements that produce speech sounds (phonemes). The classification of actual speech sounds or words yields insufficiently robust results to support a realistic technology: between 16-54% accuracy for detecting one out of a set of linguistic items (words or phoneme strings) [36]. While speech decoding from implanted electrodes succeeds at substantially more difficult classification tasks, the efficacy of such methods is boosted primarily by language prediction models, which can improve median word error detection rate up to 35% [29]. For example, words in a possible set of 50 items obtained a mean classification accuracy of 47% in post-hoc offline analyses [29]. This means that in the absence of facilitatory language prediction models, the actual speech decoding classifier performs at only slightly better than 20%, even with a data type that is more stable across sessions and possesses a significantly better signal-to-noise ratio.

Instability across sessions in the composition of the cortical activity being recorded is another key challenge to EEG speech decoding, compounded by the low signal-to-noise ratio of this data type and its propensity to generate artifacts in the process of data collection. This instability in combination with the complexity of some machine learning classifiers can require repeated recalibration of the model for each session or individual user, which is time-consuming and requires specialist knowledge. Development of an algorithm that can provide reliably accurate speech decoding across all users without frequent recalibration over time or personalization of model features for each individual user remains an area of development that has yet to be addressed in the speech decoding literature. A generalized or “universal” model of this nature would substantially aid the usability of the system for the target population outside of the research community. It is also important to note that both of the approaches reviewed here are computationally intensive and achieve their best results only offline with the benefit of large amounts of training data. For effective real-time translation that can compensate for novel words which the algorithm may not have previously encountered, an alternative to the data- and time-intensive model of traditional neural networks should be developed.

In light of these challenges, we propose that DDA can serve as a method of analysis for the reliable, quick, and accurate classification of motor EEG signals which may be adopted by a wide profile of other BCI researchers as a tool in their own research. Preliminary results suggest that TMS may aid this process by evoking a more consistent cortical-wide neural response across individuals, such that a universal DDA model for speech decoding may be developed.

## II. BACKGROUND

DDA is a powerful classification tool that combines differential embeddings with linear and nonlinear nonuniform functional delay embeddings. DDA has several advantages in comparison with CNNs, the most successful method thus far for the analysis of EEG data in speech decoding. Both CNNs and DDA effectively process high-dimensional data; however, CNNs do so by means of a high computational resources, such that they can be slow to train for complex tasks and they require a substantial amount of training data prior to performing the classification task and to avoid overfitting. Moreover, CNNs may require data augmentation when there is substantial variation within the data type to be classified. The complex architectures of CNNs require precise tuning for the problem of interest and they have difficulty modeling long distance relationships and require short data inputs to perform well. This may be one reason why simple classification tasks with one phoneme or word inputs have been performed well by CNNs, but not more complex decoding problems. Brain data is inherently high-dimensional, but the neural processes in speech decoding differ substantially in nature from the text or vision recognition tasks to which CNNs are often applied. It is widely assumed that many neural mechanisms in the brain are nonlinear [2, 27]. Thus, the integration of nonlinear dynamics into a classification analysis allows information from the data to be detected which is not observable in traditional linear methods.

Computationally, DDA boasts two key advantages over traditional machine learning analyses. Firstly, DDA requires minimal preprocessing, as it captures only the relevant dynamics for the classification problem at hand. This provides a crucial advantage, in that preprocessing may be subjective and conditioned by individual differences between participants and the artifacts incurred during data collection. Preprocessing can strip away information in the signal that is meaningful to the neural process under analysis. Eliminating this step in the pipeline also allows for a faster analysis. Secondly, DDA uses a small set of model features. In this regard, DDA has a clear advantage over CNNs. A sparse model imparts speed to the analysis, reduces the need for huge computational resources, and virtually eliminates the likelihood of overfitting the model. Figure 1 provides an overview of the two-step process by which DDA is carried out: (i) model selection, which involves choosing the model structure and parameters that best fit the overall dynamics of the EEG data. Model selection can be supervised or unsupervised. In this analysis, we take a supervised approach in order to optimize structure selection by random-subsampling cross validation, a generalization of bootstrapping. The second step involves data analysis, during which data is fitted to the chosen model. The corresponding coefficients are used as features for the classification problem.

**FIG. 1:**
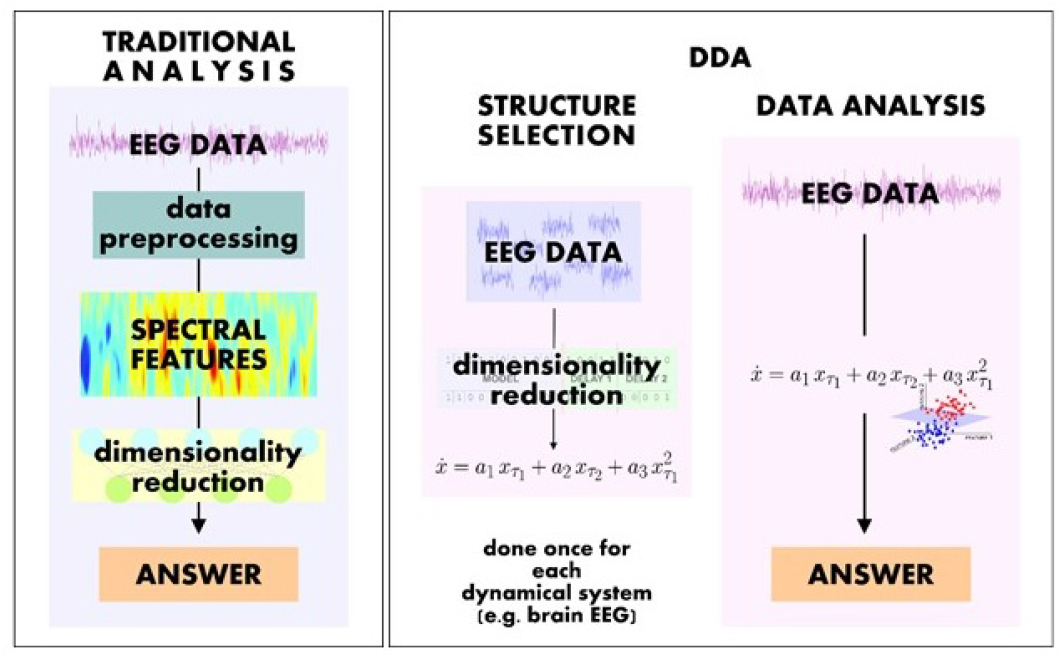
Comparison of DDA and Machine Learning Analysis.

DDA has been successfully applied to a range of data types, including dolphin echolocation data (object identification) [14], heart ECG data (differentiating heart diseases) [22, 23], EEG data (sleep staging, biomarker development for schizophrenia patients) [17, 20, 37] Parkinsonian movement data in combination with EEG data (biomarker development, disease progression assessment) [15, 16, 18, 21], and iEEG data from epilepsy patients (seizure prediction, dynamical state detection) [13, 19, 25]. In findings of particular relevance, DDA has successfully analyzed speech signals (audio signal recognition [12], emotion detection in speech [9]). This previous success illustrates the suitability of DDA for modeling neural data and physiological processes more generally.

A more exploratory approach concerns the application of TMS. TMS acts upon cortical neurons by inducing an electrical field that depolarizes the membrane potential and pushes neurons past an excitation threshold. However, there remains considerable debate as to what exact effect TMS may have upon neural processes. TMS has been argued to act upon the same neurons implicated in volitional motor commands [1], an argument that is supported by task-related TMS-EEG studies. Findings from such studies suggest that task-specific stimulation elicits task-dependent neural responses and that the functional improvements seen in task performance are related to changes in cortical reactivity found close to the stimulated region [5, 34, 39]. For example, paired pulse TMS has been shown to improve subject performance on a phoneme discrimination task when motor areas involved in the production of two categories of phonemes were stimulated: reaction times and errors in the phoneme discrimination task decreased for each phoneme when the site associated with that phoneme was targeted [7]. The logical conclusion is that stimulation of the relevant motor area leads to either (i) facilitation of the behavioral response, corresponding to an increased ability to perceive the related phoneme, due to priming of the concordant motor representation, or (ii) inhibition of the alternative behavioral response, corresponding to a reduced ability to perceive the unrelated phoneme, due to priming of the discordant motor representation.

In either case, we argue that stimulation of the motor area should therefore evoke an optimal state for detecting the relevant neural signals necessary for decoding the phoneme of interest. Generally speaking, the theoretical assumptions of this approach are commonly accepted within the speech decoding literature. Motor circuits have been shown to be involved in the speech-perception process [33] and reliance upon neural activation within areas of the motor cortex associated with speech production is standard practice for speech decoding among BCI researchers [4, 30]. The exact nature of this optimal state is less clear and remains an important question in understanding the physiological response to TMS. Discharging action potentials that are near threshold may create a homogeneous state in the stimulated region, in effect reducing the noise that typically interferes with detecting the relevant signals. In this scenario, features may be isolated that can be used as inputs for the classification analysis by means of inhibiting competing neural signals. Alternatively, TMS may augment the motor signals relevant for the decoding task. When the motor cortex is stimulated, TMS elicits motor-evoked potentials (MEPs) and cortical responses [10] termed TMS-evoked potentials (TEPs) [8], as well as a range of induced neural oscillations and connectivity changes [42, 43]. These responses are stable over sessions performed a week apart [3], indicating they could serve as inputs for machine learning classifiers. Thus, analysis of the EEG signals collected during TMS may more readily allow for identification of cortical locations with a usable signal, helping to indicate areas of focus when using EEG prospectively.

We aim to test these assumptions in performing speech decoding by means of DDA on a dataset of phonemes (4 consonants and 5 vowels) that have been targeted with stimulus-specific TMS. We propose that the combined application of TMS and DDA will allow for superior speech decoding classification by (i) optimizing the stimulus-specific information in the neural signal, and (ii) analyzing both linear and nonlinear dynamics to access the widest possible range of useful information in the neural signal.

## III. METHODS

### Experimental design

Participants listened to consonant phonemes played aurally and identified each with a button-press response. Areas in the motor cortex associated with the production of each category of phonemes were stimulated: lip and tongue regions for bilabial (/b/,/p/) and dental (/t/,/d/) phonemes, respectively (see Figure 2). To aid intelligibility, each phoneme was followed by one of five vowels (/i/,/3/α,/A/,/u/,/o℧/). Two single pulses were administered, separated by a short 50 ms interpulse interval. Pulses were timed so that the last TMS pulse occurred 50 ms prior to consonant presentation. In the stimulated trials, the two pulses were delivered at 110% of the FDI rMT. In each of 4 blocks, participants completed 80 trials, 60 with TMS and 20 random catch trials. Random unstimulated catch trials serve as a reference to evaluate the effect of TMS by indicating the baseline classification accuracy in decoding speech from EEG.

**FIG. 2:**
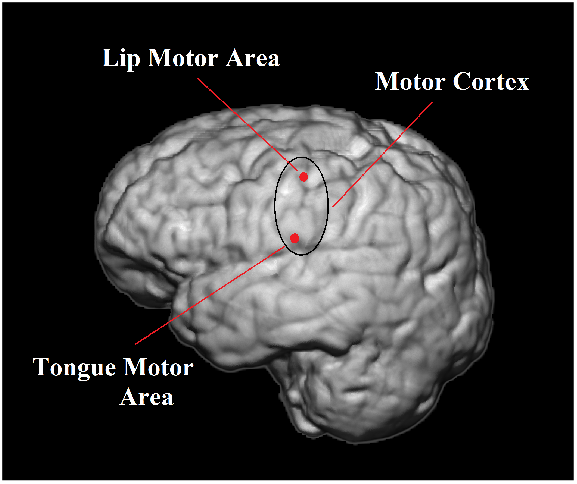
TMS Targets.

### Participants

Healthy adults between the ages of 20 and 40 were recruited by means of fliers from the UCLA community. Eligibility criteria included no prior or concurrent diagnosis of any neurological (e.g., epilepsy, Tourette’s syndrome), psychiatric (e.g., schizophrenia), or developmental (e.g., ADHD, dyslexia) disorders; and no structural brain abnormalities (e.g., aneurysm). Participants provided written informed consent and were paid for two sessions. In the first session, an MRI scan was conducted. The scan was used to target areas in the motor cortex associated with phoneme production. In the second session, participants performed a phoneme discrimination task.

### Data collection

EEG data collection was conducted in the Neuromodulation Division of the Semel Institute for Neuroscience and Human Behavior at UCLA. TMS was carried out by means of a Magstim Super Rapid Plus1 stimulator, and the stimulation targets were identified using the Visor 2 neuronavigation system (ANT Neuro). Subject-specific targets were generated for each participant by means of transforming targets previously identified in the literature for the lip and tongue motor areas [7] to the subject-space of each participant MRI scan. The TMS coil was positioned using frameless stereotaxy. Coil orientation was maintained at 45 degrees with respect to the interhemispheric fissure. Electrode positions were digitized and registered to individual subject MRIs using the ANT Neuro Xensor (ANT Neuro). EEG signals were then bandpass-filtered 0.1-350 Hz, sampled at 2000 Hz, and referenced to an additional forehead electrode. All electrode impedances were kept < 5 kΩ. All data collection measures were approved by the IRB.

### Motor thresholding

Each subject’s motor threshold was determined in the presence of a physician to select the appropriate intensity of stimulation. Motor evoked potentials (MEPs) were elicited in the first dorsal interosseus (FDI) muscle of the dominant hand at the minimum amount of stimulation needed to evoke a MEP in a hand muscle after a single pulse over the primary motor cortex. Single TMS pulses were delivered in the motor cortex contralateral to the dominant hand. The intensity of the stimulation was gradually lowered until reaching a level at which 5 out of 10 MEPs in a hand muscle have an amplitude of at least 50 microvolts.

## IV. ANALYSIS

### Delay differential analysis (DDA)

DDA combines differential embeddings with linear and nonlinear nonuniform functional delay embeddings [31, 38, 41]. The use of two delay variables relates the current derivatives of a system to current and past values of the system, [11, 13, 26]. The DDA model maps experimental data onto a set of natural embedding coordinates. The general DDA model with two delays and three terms is

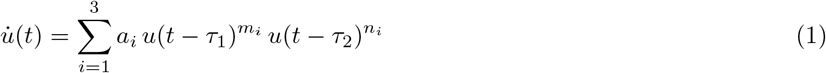

where *u*(*t*) is a time series, *m*_*i*_, *n*_*i*_, *τ*_1,2_ ∈ ℕ_0_ and a degree *m*_*j*_ + *n*_*j*_ ≤ 4 of nonlinearity. We use DDA models with two time delays and three terms to reduce complexity. Note, that the delays are independent from each other and have no physical meaning for a nonlinear DDA model [24]. They are specific to the question of research, here phoneme identification. In DDA we use the coefficients *a*_*i*_ and the least square error *ρ* as independent parameters or features. The model is fixed and is not updated during the analysis. DDA and its models have several advantages over the high dimensional feature spaces of other signal processing techniques: (i) due to sparsity of the model, the risk of overfitting is greatly reduced; (ii) it is insensitive to noise since such a tiny model concentrates on the overall dynamics of the system and cannot additionally model noise; (iii) it is computationally fast; (iv) there is no need for pre-processing except normalization to zero mean and unit variance, which will cause the model to ignore amplitude information and concentrate on system dynamics.

DDA is a two-step process. (i) For a new class of data (e.g. EEG data) the best DDA model (i.e. the coefficients *m*_*i*_ and *n*_*i*_ as well as the delays *τ*_1,2_ that best fit the overall dynamical properties of the system) need to be selected. This can be done by supervised (maximizing the classification performance) or unsupervised (minimizing the least square error) structure selection from a list of candidate models [25, 26]. The performance is evaluated by the area under the receiver operating characteristic (ROC) curve (AUC or A’). This is essentially a plot of the true positive rate against the false positive rate. This step is done once and does not have to be repeated for new data from the same data class (e.g. EEG data). For the question of interest (here phoneme classification), the best delays were selected. There is one model that has been found to represent the overall nonlinear dynamical structure of EEG data:

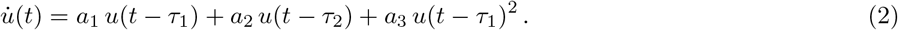

This model appears unique to EEG data [25]. (ii) After the DDA model is established, data can be analyzed by fitting the data to the model and estimating the features (*a*_1_, *a*_2_, *a*_3_, *ρ*). We estimate the parameters from the data without any pre-processing or filtering except normalizing each data window to zero mean and unit variance.

Prior to conducting the classification analysis, the EEG data was spliced into individual items (consonant/vowel pairs) of roughly 250 ms in length. A 35 ms window from consonant onset was used to ensure that the auditory input from vowels did not contaminate the consonant classification problem. While alternative approaches may utilize a larger time window in order to capture the full range of evoked auditory potentials, this decision has a precedent in previous literature that found a short time window at phoneme onset to be the most informative for speech decoding [30].

In the current study, we identify one out of four possible consonant phonemes by means of a four-way classification in which the best delays *τ*_1,2_ are selected for each consonant to be distinguished from the other three phonemes. The targets for the four classifiers C_1,2,3,4_ and the four phonemes are shown in Table I. We applied all four classifiers using a one-against-all approach in which each phoneme is treated as the positive class, one at a time. The classifier with the highest value is selected as the correct label. Performance is reported as the percentage of correctly identified phonemes.

**TABLE I:** Phoneme targets for the four-way classifiers.

|  | /b/ | /p/ | /t/ | /d/ |
| --- | --- | --- | --- | --- |
| $C_1$ | 1 | 0 | 0 | 0 |
| $C_2$ | 0 | 1 | 0 | 0 |
| $C_3$ | 0 | 0 | 1 | 0 |
| $C_4$ | 0 | 0 | 0 | 1 |

An exact comparison of our method and that of previous authors, which would account for all possible experimental design factors, is not possible due to a lack of access to the data sets used by other research teams. However, the percentage of correctly identified phonemes (accuracy) is a common metric in speech decoding papers that can be approximately interpreted across studies. Precision and recall metrics are also frequently reported. Precision is the proportion of classified phonemes that match the correct label, whereas recall is the proportion of phonemes that were detected in the speech stream. While the difference between these two metrics is interesting for theoretical reasons [6], overall accuracy is more relevant concern for the effectiveness of a classification analysis in real world terms. Therefore, in this short report we will indicate classification accuracy and use this metric as an estimate for how analyses performed by different algorithms on different data sets can be evaluated.

## V. RESULTS

This study hypothesized that TMS would (i) improve the classification performance for the corresponding TMS target/phoneme pairs, and (ii) allow for better classification across participants. Two approaches were evaluated: a) individual models, in which model parameters were optimized for each participant under each target condition (sham, lip TMS, or tongue TMS) and for each channel; and b) a generalized model, in which the best overall model parameters were selected across participants and channels for each condition. Data comprised approximately 20 trials per condition, participant, and channel.

### Individual models

We first identified the optimal model using cross-validation and tested this model on all available data. The best model parameters were selected on all trials for each condition, participant, and channel (80 total) using a 56/24-fold repeated random subsampling cross-validation, repeated 20 times for a one-vs-all classifier for each of the 4 phonemes. This resulted in 6 (participants) * 3 (conditions) * 61 (channels) * 4 (phonemes) = 4392 classifiers.

Figure 3 shows modified confusion matrices in the top three rows that correspond to the three conditions and six subjects and the accuracy for each phoneme is summarized in the bottom row. The results illustrate that DDA can achieve essentially perfect classification. Moreover, perfect separation could be found with just one EEG channel, optimized for each sound and participant. Small data sets may be at risk of overfitting; however, with our sparse 3-term model, 20 trials should be sufficient and should not lead to overfitting. In the second analysis, we wanted to confirm this result in a second, more stringent classification task. Because we only had around 20 trials per subject, channel, and condition, we selected the best model on 5 trials from one channel and added 5 trials from a neighboring channel for structure selection. We then tested on the leave-out data from each channel separately.

**FIG. 3:**
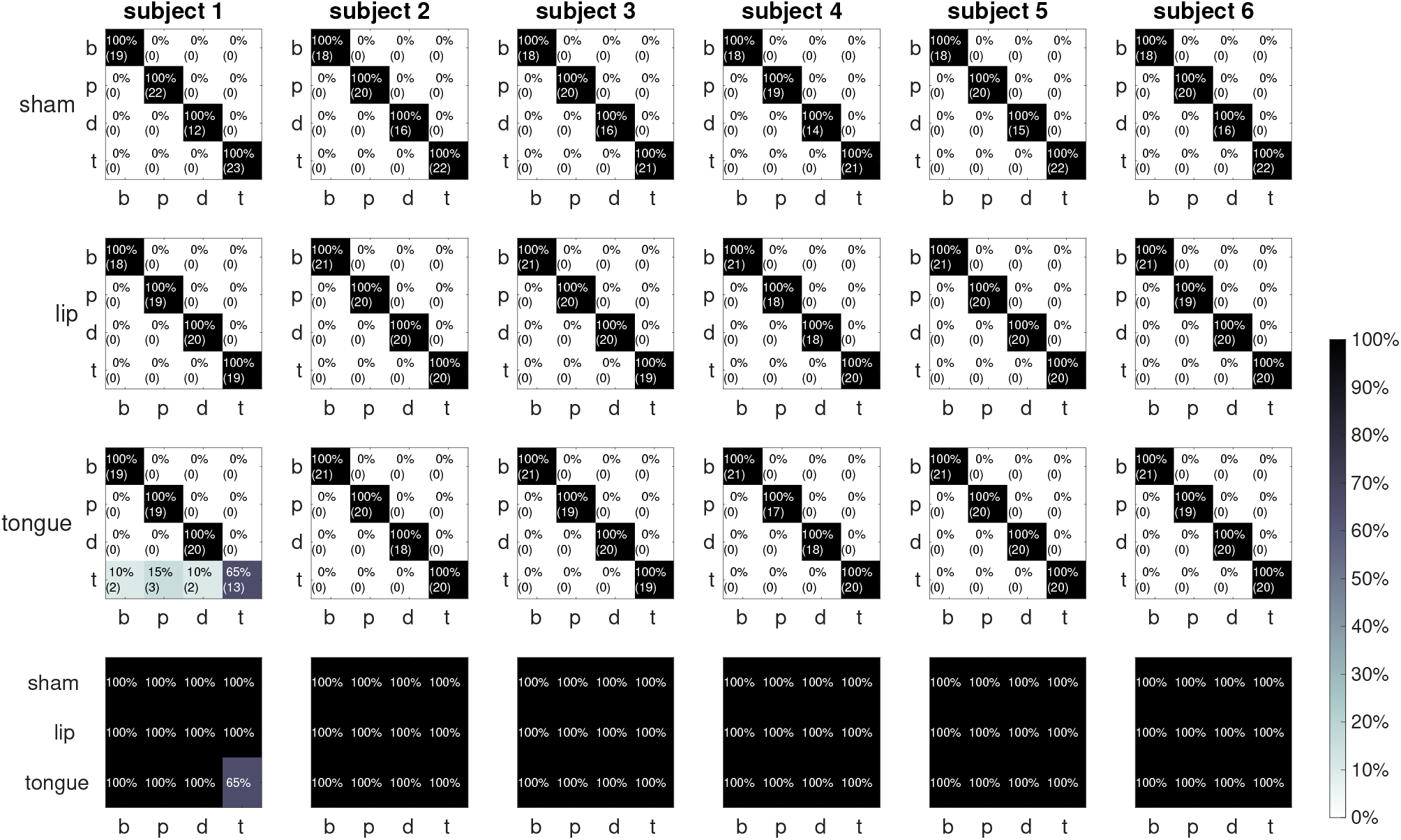
Classification on all data for individual models for each subject, channel, and condition.

Fig. 4 shows modified confusion matrices in the three top rows and the accuracy for each phoneme in the bottom row. We observe the potential facilitation of classification by TMS, although the sham condition also performs well in select participants. At this stage, we cannot state with certainty whether the hypotheses regarding TMS target/phoneme pairs are upheld. TMS was also theorized to increase the similarity in cortical response across participants. The possible effect of TMS will become more apparent when a generalized model is assessed. Already at this stage of the analysis, significant improvement in classification accuracy relative previous EEG speech decoding studies is observed.

**FIG. 4:**
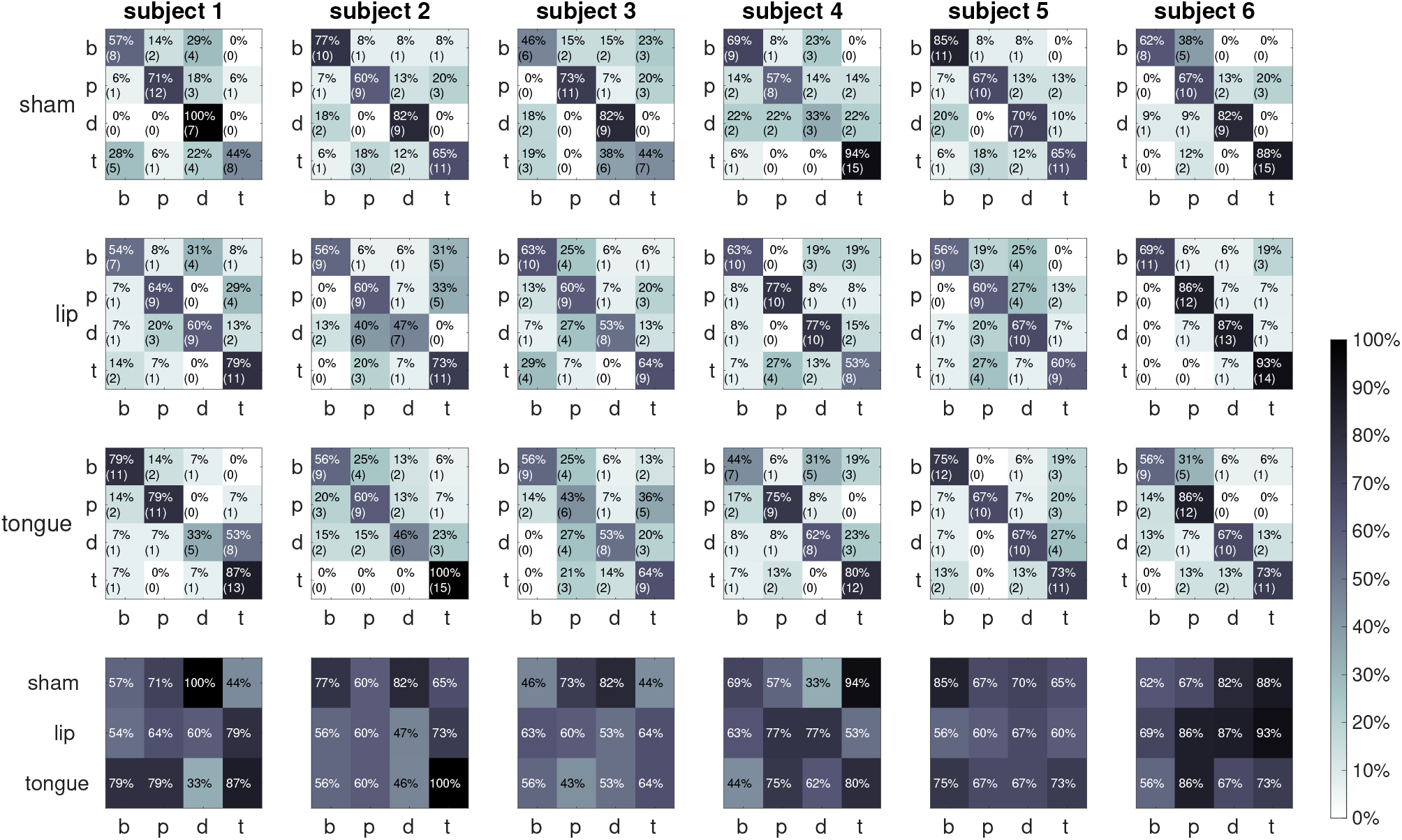
Classification on leave-out data for individual models for each subject, channel, and condition.

### Generalized model

The best model parameters were selected on 10 trials from each condition across channels and subjects for each condition and phoneme. In this case, there are four general classifiers (four phonemes) for each condition, but the best channel location depends on the subject. We find that in this more generalized approach, without individualized selection of all model parameters, superior classification is obtained: >71% correct for each phoneme in at least one of the three conditions. In many cases, classification reaches 100% accuracy (see Fig. 5). It is notable that a more generalized approach appears to outperform the fully individualized models. However, this analysis was also performed on a larger data set from all six subjects.

**FIG. 5:**
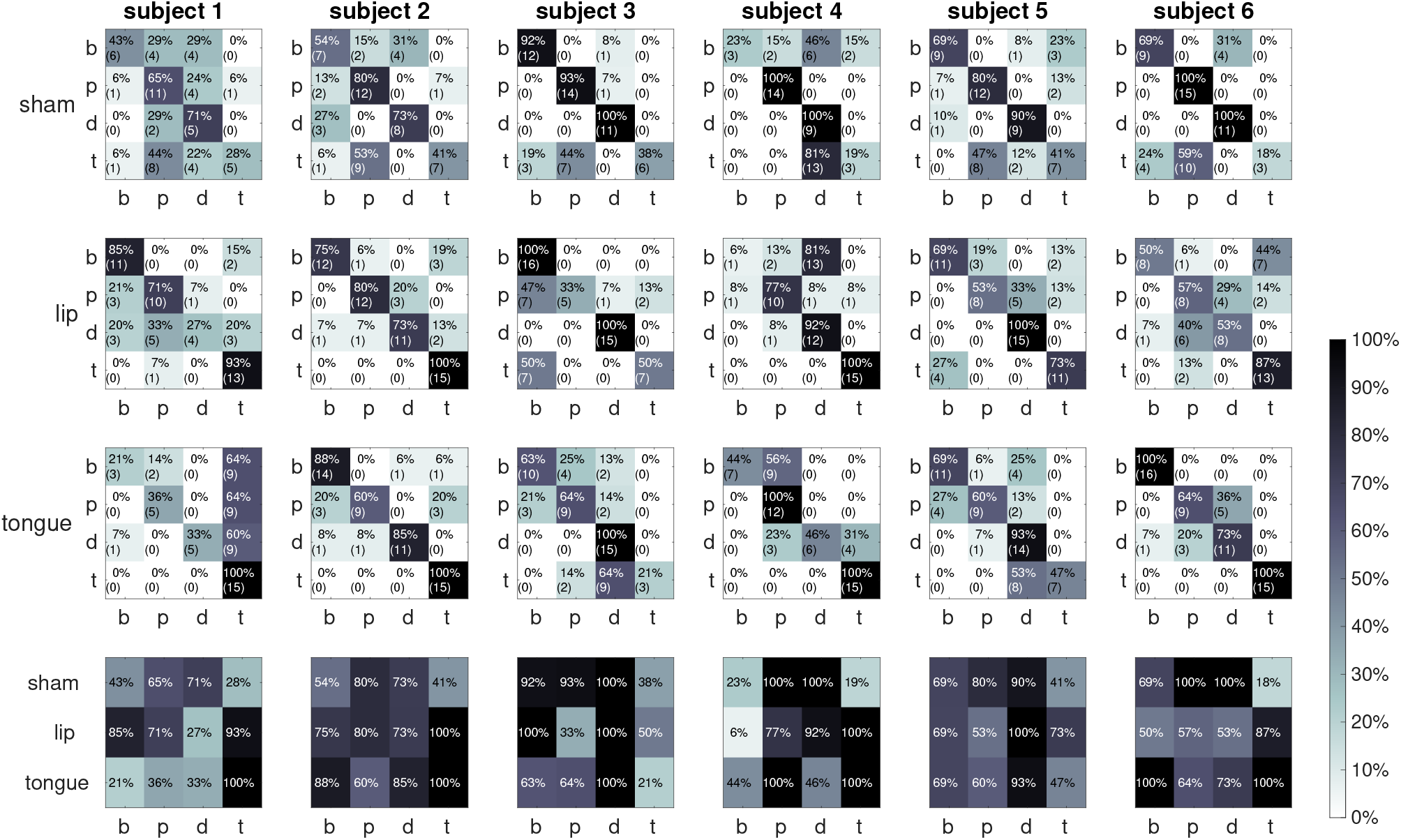
Classification on leave-out data for generalized models for each condition and phoneme.

The two TMS conditions taken together tend to provide better classification accuracy than in the sham condition in most participants. This may support the ability of TMS to augment or isolate the neural signal of interest in EEG signals. Given the substantial variation in results across participants, more data is needed to better understand this effect. We argue that discrepancies between participants and in regard to our hypotheses may reflect variation in the optimal target location for each participant: the target coordinates were taken from the literature and transformed to subject-space during the neuronavigation targeting process. Nonetheless, considerable differences in the functional organization of the brain are found between individuals. Targets selected as the average coordinates of multiple subjects may therefore not reflect the cortical regions underlying motor representations for each individual with the same degree of accuracy.

However, when we observe the minimum of the four classifiers we find greater support that TMS may create a more uniform neural response to phonemes across subjects. In this analysis, we identified when a phoneme could not be detected. The best classification performance on the leave-out data for each subject, condition, and channel is shown in Figure 6. Plots represent the signal obtained by each electrode. Each plot consists of a matrix of 24 cells: the six rows are the results for each of the six subjects across the 4 phonemes, shown in the columns. The results show clear suppression of /d/ in the lip TMS condition and of /b/ is the tongue TMS condition (i.e., the phoneme not associated with the target). There is greater consistency in the classification results obtained across participants and across all electrode channels in both of the TMS conditions than in the sham condition. Consonants with the articulatory feature of ‘voicing’ (vocal cord vibration) - /b/ and /d/ - show the greatest effect. Voicing requires greater motor activity, and thus this result supports our claim that TMS targets the motor cortex associated with phoneme articulation.

**FIG. 6:**
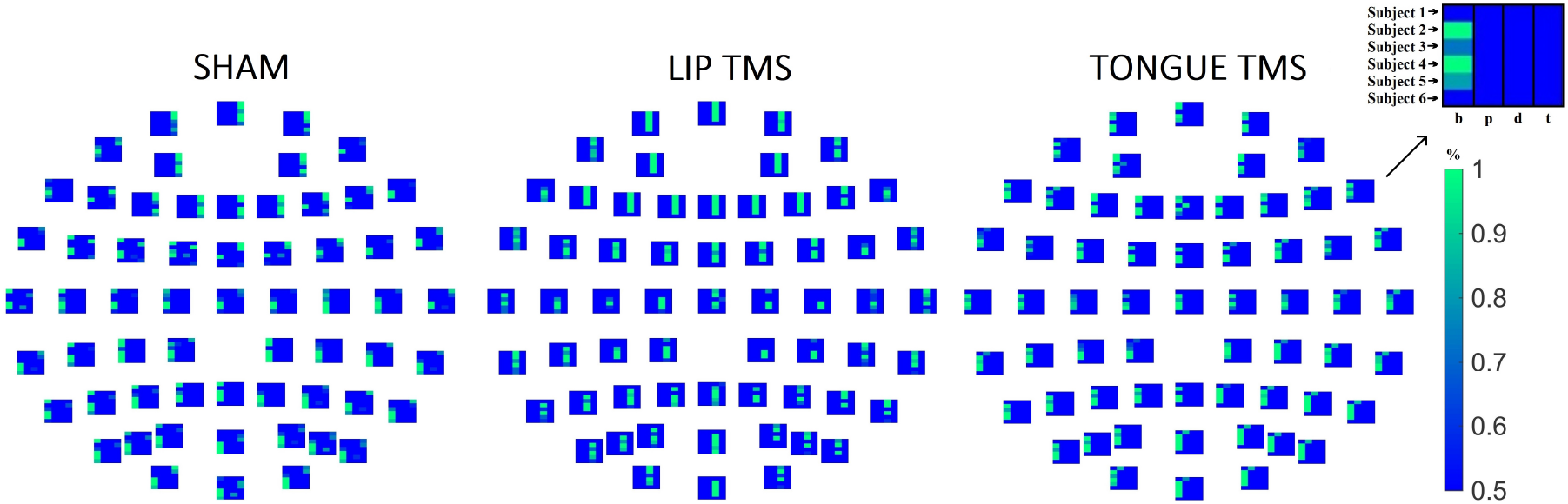
Classification on leave-out data per electrode channel. Generalized model.

## VI. CONCLUSION

While preliminary in scope, our results clearly indicate the superior capabilities of DDA for speech decoding. Even with the low signal-to-noise ratio of EEG data, the analysis achieved an astonishing 70-100% classification accuracy for all phonemes in at least one of the three conditions when tested on leave-out data. These results have a clear advantage over those obtained by previous EEG speech decoding studies (20%-54%). Moreover, DDA more than doubles classification accuracy while performing a more complex classification task with a minimal data set: in the first two analyses, superior performance was achieved with data from just one EEG channel. DDA even outperformed the metrics reported for studies utilizing invasive data (47%). In this case, although our analysis is less complex, it is performed without the added assistance of predictive language models. There also appears to be good evidence that TMS may assist in creating a consistent cortical response across participants, which in turn may facilitate the development of a generalized model. In select participants, TMS promoted greater classification accuracy. Additional research into subject-specific TMS coordinates will allow for the better understanding of how to harness the effects of TMS for speech decoding.

## Notes

### Competing Interest Statement

The authors have declared no competing interest.

